# Cold hardiness-informed budbreak reveals role of freezing temperatures and daily fluctuation in chill accumulation model

**DOI:** 10.1101/2024.02.25.581952

**Authors:** Michael G. North, Beth Ann Workmaster, Amaya Atucha, Al P. Kovaleski

**Affiliations:** Department of Plant and Agroecosystem Sciences, University of Wisconsin–Madison, Madison, WI 53706, USA

**Keywords:** budbreak, chill accumulation, chill model, cold hardiness, daily amplitude, dormancy, freezing, temperature

## Abstract

Fundamental questions in bud dormancy remain, including what temperatures fulfill dormancy requirements (i.e., chill accumulation). Recent studies demonstrate freezing temperatures promote chill accumulation and cold hardiness influences time to budbreak – the phenotype used for dormancy evaluations. Here we evaluated bud cold hardiness (*CH*) and budbreak responses of grapevines (*Vitis* hybrids) throughout chill accumulation under three treatments: constant (5°C), fluctuating (−3.5 to 6.5 °C daily), and field conditions (Madison, WI, USA). Chill treatments experiencing lower temperatures promoted greater gains in cold hardiness (*CH*_field_>*CH*_fluctuating_>*CH*_constant_). All treatments decreased observed time to budbreak with increased chill accumulation. However, perceived treatment effectiveness changed when time to budbreak was adjusted to remove cold acclimation effects. Among three classic chill models (North Carolina, Utah, and Dynamic), none were able to correctly describe adjusted time to budbreak responses to chill accumulation. Thus, a new model is proposed that expands the range of chill accumulation temperatures to include freezing temperatures and enhances chill accumulation under fluctuating temperature conditions. Most importantly, our analysis demonstrates adjustments for uneven acclimation change the perceived effectiveness of chill treatments. Therefore, future work in bud dormancy would benefit from simultaneously evaluating cold hardiness.

**Highlight:** A new chill accumulation model demonstrates how bud cold hardiness changes elicited by chill treatments affect the interpretation of thermal effectiveness in promoting dormancy progression and release.

## Introduction

Chilling – the cumulative exposure to low temperatures – is essential for woody perennial plants to harmonize cycles between dormancy and growth with the annual rhythm of seasons, particularly in temperate and boreal climates (Chuine and Régnière, 2017; Lang, 1987, Weiser, 1970). Once a plant enters a state of dormancy, growth ceases and metabolic activity decreases until enough time is spent at chilling temperatures to fulfill a certain chilling requirement and plants overcome mechanisms preventing growth (Samish, 1954; Faust et al., 1997; Campoy et al., 2011). These important seasonal cycles have been the focus of a long tradition of studies investigating temperature’s role in dormancy regulation (Knight, 1801), including the development of chill models, which generate low temperature-related thermal time units to quantify chill accumulation and delineate dormancy status (Coville, 1920; Vegis, 1964; Fuchigami et al., 1982; Saure, 1985; Shaltout and Unrath, 1983; Richardson et al., 1974; Fishman et al., 1987a,b). Chill models are not based on a functional understanding of physiology or genetics, as those are currently too limited (Cooke et al., 2012; Fadon et al., 2020), and instead approximate this largely unknown biological process based on correlating environmental conditions (Luedeling, 2012). Chill models that are widely used today still require local adaptation (Luedeling and Brown, 2011), indicating they do not accurately describe the relationship between temperature and dormancy progression. Therefore, progress in dormancy research, from molecular level to ecosystem adaptation to future climates, still requires a better understanding of how temperatures promote chill accumulation.

Dormancy is typically evaluated based on time to budbreak (or related visible growth metrics, such as percent budbreak within a period of time) after exposure to chilling (Coville, 1920; Londo and Johnson, 2014; Baumgarten et al., 2021). In experiments, time to budbreak [sometimes expressed as heat requirement, using growing degree-days or equivalent (e.g., Pope et al. (2014)] decreases with longer exposure to chilling treatments. However, the rate of change in budbreak responses varies depending on the temperature in chilling treatments (Fig 1A) since not all temperatures have the same magnitude of effect toward overcoming the mechanisms preventing bud growth (i.e., chilling efficiency). Chill models are the tools used to describe the different chilling efficiencies of temperatures and they do so by integrating an assigned temperature-specific efficiency and time into thermal time units [Fig 1B; e.g., Shaltout and Unrath (1983); Richardson et al. (1974); Fishman et al. (1987a, b)]. If a chill model incorrectly describes the relative chilling efficiency across a range of temperatures, then the budbreak response to chilling treatments at different temperatures will still be distinct when characterized according to the model’s thermal time units (Fig 1C). However, if a chill model is correctly describing temperature-chill efficiency, the budbreak response between treatments should be indiscernible (Fig 1D). In theory, a correct model such as this would describe budbreak responses similarly across different locations and/or in varying experimental chill treatments for a particular genotype.

**Figure 1.**
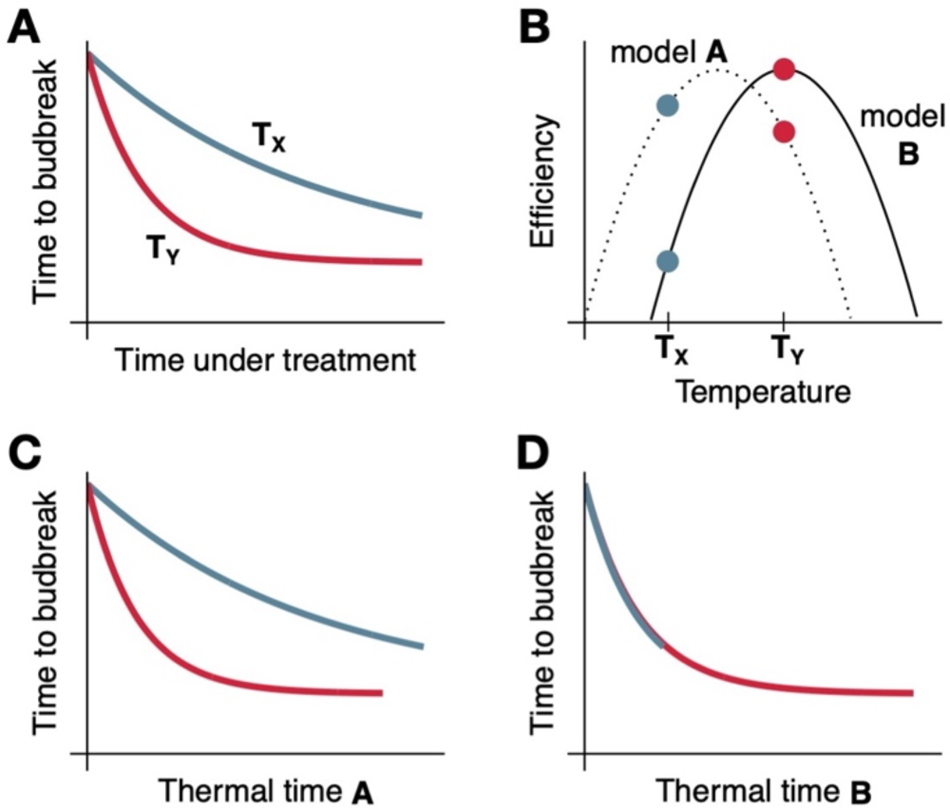
Simplified conceptual method of evaluation for chill accumulation models though budbreak assays. **(A)** Time to budbreak assays are conducted where buds are exposed to different temperatures (e.g., here T_X_ and T_Y_, where T_Y_ > T_X_) for certain amounts of time to accumulate chill, then placed under forcing conditions (i.e., warm temperatures, long days) to evaluate time to budbreak. **(B)** Different chill models assign different efficiency levels for temperatures – for model A, efficiency of T_X_ is greater than T_Y_, whereas the opposite is true for model B. Time is then converted into thermal time (i.e., chill units) based on temperature efficiencies **(C, D)**. Where poor descriptions of temperature efficiencies are built into the model, differences between treatment temperatures that have the same chill unit accumulation can separate further than when ordinary time is used **(C)**. When temperature efficiencies are correctly described, treatments collapse into a single line, as the differences provenient from temperatures are accounted for **(D)**.

The majority of chill models are “time-homogeneous”: an hour spent at a certain temperature will always provide the same number of chill units (Zhang and Taylor, 2011). However, several studies have shown that chill treatments with fluctuating temperatures can enhance efficiency for a given temperature (Couvillon and Erez, 1985; Anzanello et al., 2014; Horikoshi et al., 2017). This thus requires a “time-inhomogeneous” model, where factors surrounding the time when a certain temperature is experienced affects how many chill units are accumulated. One widely used chill model, the Dynamic Model, directly integrates an enhancement in chill unit accumulation based on the sequence of temperatures experienced, (Fishman et al., 1987a, b; Erez et al., 1990). However, despite the speculated mechanistic basis for the Dynamic Model, it still fails to accurately describe chill accumulation correctly across different environments (Luedeling, 2012). Another overlooked factor in current chill models is the incorporation of freezing temperatures in the chill accumulation temperature range, even when a growing body of literature suggests freezing temperatures should be included in the chilling range (Mahmood et al., 2000; Rose and Cameron, 2009; Guak and Neilsen, 2013; Cragin et al., 2017; Baumgarten et al., 2021).

Several recent studies have shown time to budbreak is significantly influenced by bud cold hardiness, which refers to the minimum freezing temperature buds can survive exposure to (lower temperature values mean more cold hardy buds) (Fig. 2) (Kovaleski et al., 2018; Kovaleski, 2022; North et al., 2022). Cold hardiness is a dynamic trait that changes throughout the dormant season in the field, largely because of temperatures experienced (Weiser, 1970). It has been posited that exposure to chill treatment temperatures may promote changes in cold hardiness, in addition to progression of dormancy, that could result in earlier or later budbreak depending on direction and magnitude of cold hardiness changes (Fig. 2). This combined effect of cold hardiness and chill accumulation has been evaluated in field-chilled material (Kovaleski et al., 2018; North et al., 2022; Kovaleski 2022). However, artificial chilling treatments are typically not as cold as most temperate environments, even when considering studies that have included freezing temperatures (Mahmood et al., 2000; Rose and Cameron, 2009; Guak and Neilsen, 2013; Cragin et al., 2017; Baumgarten et al., 2021). Based on cold hardiness models, warmer temperatures lead to smaller rates of cold hardiness gain (Ferguson et al., 2011, 2014; North et al., 2022; Kovaleski et al., 2023; Jones et al., 2023), but can also not elicit a genotypes’ full potential in terms of cold hardiness (Kovaleski et al., 2023; Jones et al., 2023). Thus, different artificial chilling treatments likely promote uneven levels of cold acclimation.

**Figure 2.**
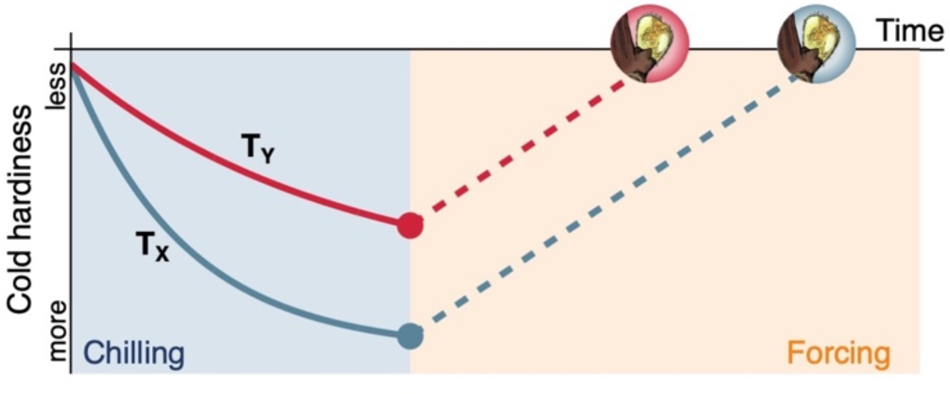
Potential effects of chilling treatments on cold hardiness and time to budbreak. During chilling, samples can gain cold hardiness to different levels, depending on chilling temperature (e.g., here T_X_ and T_Y_, where T_Y_ > T_X_). Once samples are moved into equal and constant forcing conditions (i.e., warm conditions where T_forcing_>T_Y_>T_X_), budbreak occurs after most cold hardiness is lost. Therefore, more cold hardy samples take longer to break bud when the deacclimation rate is the same [many other scenarios related to cold hardiness dynamics are explored within North and Kovaleski (2024)].

Chill model estimations of chilling requirements are not generalizable across geographic locations even for clonally propagated plants (Luedeling and Brown, 2011). These shortcomings limit the practical utility of chill models, but also compromise dormancy evaluations for mechanistic studies in physiology and genetics. We hypothesize that discrepancies in budbreak responses in studies that span different natural chill locations or use different artificial chill treatments are associated with changes in bud cold hardiness. Here, we use hybrid grapevines (*Vitis* interspecific hybrids) as a model species under natural and artificial chilling conditions to (1) evaluate the magnitude of cold hardiness changes related to chill treatments and (2) incorporate cold hardiness changes into our budbreak observations for the evaluation of chill models.

## Materials and Methods

### Site, plant material, and sampling

This study was conducted throughout three winter seasons, 2018-2019, 2019-2020, and 2021-2022, using plant material from vineyards at the West Madison Agricultural Research Station (WMARS) in Verona, WI, USA (43° 03’ 37” N, 89° 31’ 54” W). All experiments throughout this study included bud material from five interspecific hybrid grapevine (*Vitis* hybrid) cultivars: ‘Brianna’, ‘Frontenac’, ‘La Crescent’, ‘Marquette’, and ‘Petite Pearl’. Buds from node positions three to ten were sampled from healthy canes that grew in the previous growing season. Canes with buds were collected in plastic bags, stored on ice during transportation, and prepared immediately upon arrival at the laboratory.

### Chill treatments

Buds were conditioned in three chilling treatments: (1) “field”, (2) “constant” and (3) “fluctuating”. For field treatment, buds were exposed to ambient air temperatures in the vineyard as intact canes. For the constant treatment, buds were maintained at a constant 5±0.5°C. For the fluctuating chill treatment, buds were exposed to a 24-hour repeating cycle that included 6 hours at −3.5±0.5 °C and 16 hours at 6.5±0.5 °C with one-hour ramps between the two set temperatures.

For the constant and fluctuating chill treatments, buds were sampled in mid-October of each year (11 October 2018; 15 October 2019; 13 October 2021). Cane cuttings were randomized across vines and blocks, wrapped in bundles at either end with moist paper towels, then sealed in plastic bags. The bagged cuttings were placed in dark temperature chambers programmed for the constant and fluctuating chill treatments. For the field treatment, canes were collected from the vineyard on approximately the same dates as sets of cuttings were subsampled from the constant and fluctuating treatments. The field treatment was sampled throughout the entire dormant season (≥ 21 weeks, depending on season), whereas artificial chilling was provided for a maximum of 14 weeks.

### Bud forcing assays

At approximately bi-weekly intervals, a subsample of canes was removed from each chill treatment for bud forcing assays. Canes were cut into single-node cuttings and placed in square pots arranged in plastic seedling trays filled with deionized water [n=15 (2021-22) or n=25 (2018-19, 2019-20) per cultivar from each treatment]. The trays with cuttings were placed under forcing conditions at 22±1.5 °C with a 16-hour photoperiod. The growth stage of the bud on each single-node cutting was evaluated daily for up to 70 days from the date of collection, and the date of budbreak was recorded. Budbreak was defined as Stage 3 (wooly bud) of the modified E-L system (Coombe, 1995).

### Cold hardiness evaluation

When each subsample was removed for bud forcing assays, a second subset of canes was also used for cold hardiness evaluation. Individual buds (15 to 30 buds per cultivar from each treatment) were excised from canes to estimate their low temperature exotherms (LTE) – the killing point temperature – with differential thermal analysis (DTA) (Londo et al., 2023; Mills et al., 2006; North et al., 2021). The equipment used includes thermoelectric modules (TEMs) (model HP-127-1.4-1.5-74 and model SP-254-1.0-1.3, TE Technology, Traverse City, MI, USA) to detect exothermic freezing reactions and a copper-constantan (Type T) thermocouple (22 AWG) or thermistor (Type K) positioned in proximity to the TEM units to monitor temperature. Trays with TEMs and thermocouples were loaded in a Tenney Model T2C or Tenney Model TJC programmable freezing chamber (Thermal Product Solutions, New Columbia, PA) connected to a Keithley 2700-DAQ-40 or Keithley DAQ6510 multimeter data acquisition system (Keithley Instruments, Cleveland, OH). Slight differences were present in the cooling protocol in the third season compared to the previous two, which do not affect measurements (Londo et al., 2023). Specifically, in 2018-19 and 2019-20, the trays were cooled to 4 °C and conditioned for one hour, then cooled to 0 °C and conditioned for one hour, then cooled to −44 °C at a rate of 4 °C per hour. In 2021-22, trays were cooled to −5 °C and conditioned for five hours, then cooled to −55 °C at a rate of 6 °C per hour.

### Adjusted budbreak based on a cold hardiness correction

To account for differences in initial cold hardiness across forcing assays, time to budbreak observations were systematically adjusted based on cold hardiness at the initiation of forcing assays. First, cold hardiness was estimated with DTA in tandem with establishing forcing assays for the field chill treatment in all seasons and for the constant and fluctuating chill treatments in 2021-22. For artificial chill treatments, cold hardiness in 2018-19 and 2019-20 was estimated based on a logarithmic model of each artificial treatment’s 2021-22 cold hardiness response. Second, using the initial cold hardiness that was estimated simultaneously with each forcing assay, a cold hardiness-based time correction was calculated following the formula:

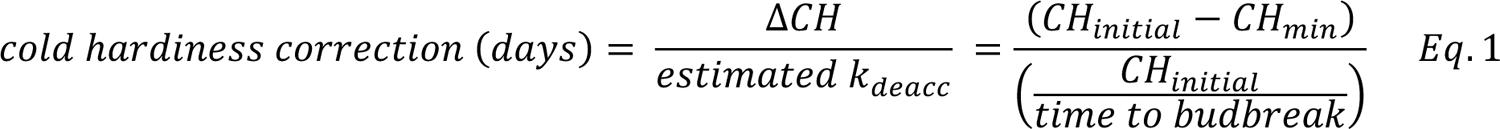

where CH_initial_ (°C) is the cold hardiness at the start of each forcing assay, CH_min_ (°C) is the lowest cold hardiness measured at the outset of chilling treatments (e.g., cold hardiness in early October, when buds were the least cold hardy), and time to budbreak (days) is the observed time to budbreak for each bud in forcing conditions. CH_initial_ varies by cultivar, chilling treatment, and time under treatment. Whereas CH_min_ was a fixed term for all cultivars and all years based on the least cold hardy point that was measured (−10.6 °C, cultivar La Crescent in season 2018-19). The denominator (CH_initial_/time to budbreak) is analogous to a deacclimation rate (unit is °C/day).

Finally, an adjusted budbreak was calculated by subtracting the cold hardiness correction from the observed budbreak (i.e., adjusted budbreak = observed budbreak – cold hardiness correction).

### Chill unit calculation

Hourly temperature data were used to calculate chill units according to several existing models in R (ver. 4.3.1; R Core Team 2023). For field conditions, hourly weather data for all three seasons were retrieved from Network for Environment and Weather Applications (NEWA; http://www.newa.cornell.edu/) participating station (Model MK-III SP running IP-100 software; Rainwise, Trenton, ME, USA) located at WMARS (see weather station “Verona (West Mad Ag Sta), WI”). Temperature in the artificial chilling treatments was monitored using a temperature logger (HOBO Model MX2305, Onset, Bourne, MA, USA). The three classic chill models used in this study were: the North Carolina Model (“NC”; Shaltout and Unrath, 1983), the Utah Model (“UT”; Richardson et al., 1974), and the Dynamic Model (“DM”; Fishman et al., 1987 a, b). A restricted cubic spline was fit with the “rms package” (Harrell 2023) to calculate chill units for the NC model. The “chillR” package (Luedeling et al., 2023) was used to calculate chill units for the UT and DM models. Because there are negative chill units in the UT and NC models, cumulative chill was calculated in two steps to determine the start date for each season. First, the cumulative chill was calculated starting from September 1. Then, the date with the most negative chill was identified and cumulative chill was re-calculated with this as the start date (i.e., new date for 0 chill unit accumulation).

We also created a novel chill model that calculates hourly chill units in a two-step process (from here on referred to as “New Model”). The new model was created to possibly incorporate a wider range of temperatures that provide chilling (i.e., negative temperatures), and to enhance chilling according to daily temperature fluctuations. First, a chill unit temperature response curve based on a temperature-photosynthesis curve (O’Neil et al. 1972) with three parameters was fit:

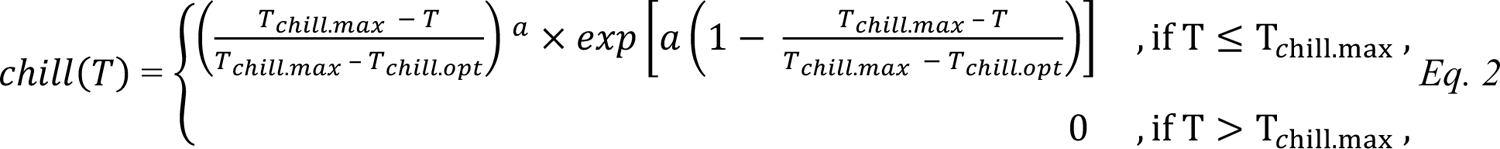

where T_chill.max_ (°C) is the maximum temperature for chilling, T_chill.opt_ (°C) is the optimum temperature for chilling, *“a”* (no units) is the coefficient related to the slope of the curve, and T (°C) is the hourly ambient temperature.

Second, a daily chill enhancement multiplying factor based on temperature amplitude was fit using a three-parameter logistic:

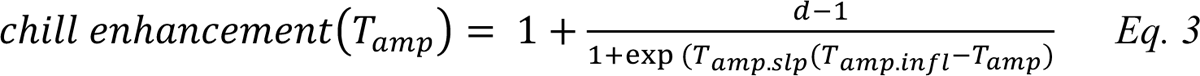

where T_amp.infl_ (°C) is the inflection point of the curve, T_amp.slp_ (no units) is the slope of the curve, “d” (no units) is the upper chill enhancement limit, and T_amp_ (°C) is the daily temperature amplitude (T_max_ – T_min_).

The final New Model calculation included the hourly chill value multiplied by the chill enhancement based on the corresponding daily temperature amplitude:

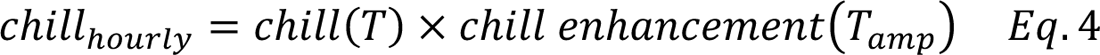

which was then cumulatively summed across the duration of chill treatments. A stepwise iterative method tested 313,920 parameter combinations for the new chill model (Table S1). Briefly, the parameter combinations included iterations where negative temperatures were both included and excluded from the chill efficiency curve, and iterations where temperature fluctuations did not enhance accumulation (chill enhancement = 1) or enhanced chill accumulation at a maximum of 1.5- or 2-fold. To evaluate the New Model, an nls() model was fit according the framework presented by Kovaleski (2022), where there are three sources of variation in time to budbreak (i.e., 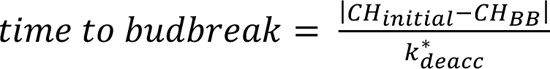). The parameter combination that minimized the RMSE between predicted and observed average budbreak (pooled across all cultivars and years) was selected for subsequent analysis.

### Chill model comparisons

To statistically evaluate suitability of each chill model for describing chill accumulation as a result of treatments, an exponentially declining curve (Cannell and Smith, 1983; Pope et al., 2014, Darbyshire et al., 2016) was fit using nls(),

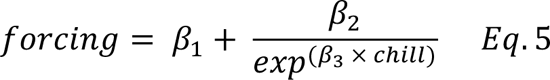

where: *forcing* is both mean observed and mean adjusted time to budbreak at 22 °C (days), *chill* is the amount of chill accumulated, β_1_ is the lower asymptote of time to budbreak at high levels of chill accumulation, β_2_ is the difference between the lower asymptote and the time to budbreak at very low levels of chill accumulation (i.e., β_1_+β_2_= intercept) and β_3_ is the slope associated with the decrease in time to budbreak with chilling accumulation. RMSE was calculated for each chill treatment separately and for chill treatments pooled together based on a model fit for all data combined.

## Results

### Cold hardiness changes during chilling

All the chill treatments elicited changes in bud cold hardiness but the responses varied in magnitude depending on the treatment. Buds in the field treatment gained the most cold hardiness (before loss in the spring), followed by the fluctuating treatment, and lastly the constant treatment (Fig. 3). The largest gains in cold hardiness occurred within the first six weeks for all treatments. Field samples continued to gain cold hardiness as temperatures decrease seasonally and lose cold hardiness (or deacclimate) upon return of warmer temperatures in spring (Fig. S1).

**Figure 3.**
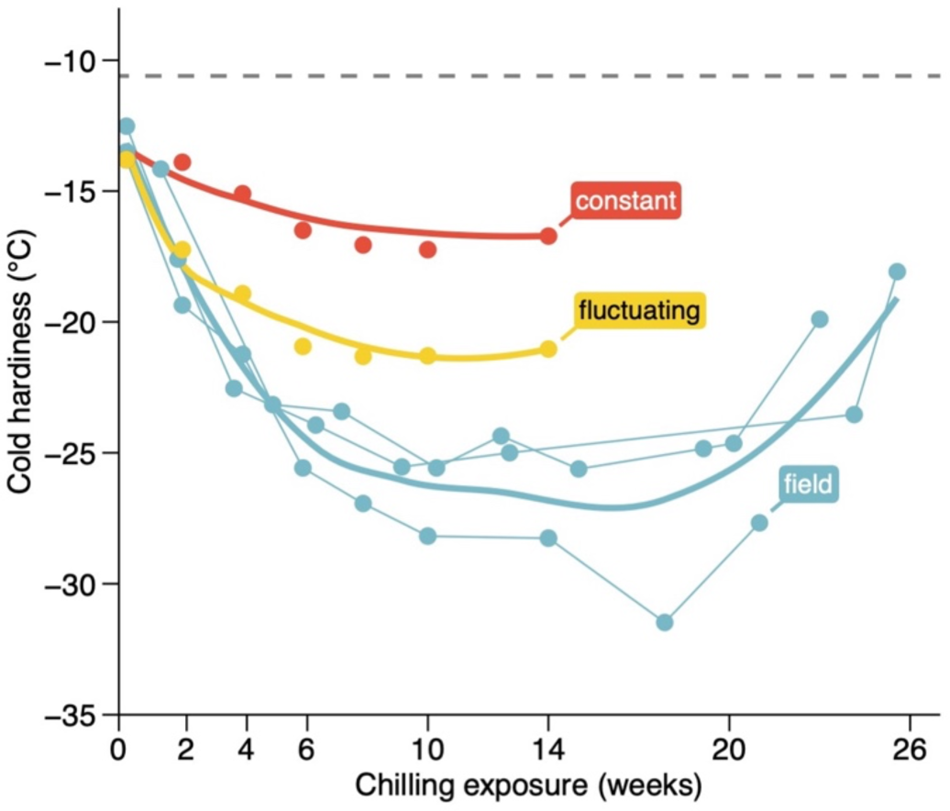
Cold hardiness changes of grapevine buds during chilling. Cold hardiness of buds of interspecific hybrid grapevines (*Vitis* hybrid) over time, chilled at three different temperatures: constant (5°C), fluctuating (−3.5 to 6.5°C daily), and field conditions in Madison, WI, USA. Dashed line indicates minimum cold hardiness measured (–10.6 °C, for *Vitis* hybrid ‘La Crescent’ cultivar in season 2018-19; CH_min_ in Eq. 1), used for cold hardiness adjustment to budbreak. Thin lines connect data points for each field season. Bold lines are loess fits of cold hardiness for each treatment in response to time.

### Chill accumulation for different models

Based on the three classic chill models, chill accumulation occurred at different rates for each temperature treatment. However, the rank order of each treatment within a model was the same: constant, followed by fluctuating, and then field. (Fig. 4). In fact, the field treatment, despite being longer in time, still accumulated less units using the NC and UT models over the course of an entire dormant season (≥ 21 weeks) compared to 14 weeks in the artificial chilling treatments (Fig 4 for 2021-22, Fig. S2 for all seasons).

**Figure 4.**
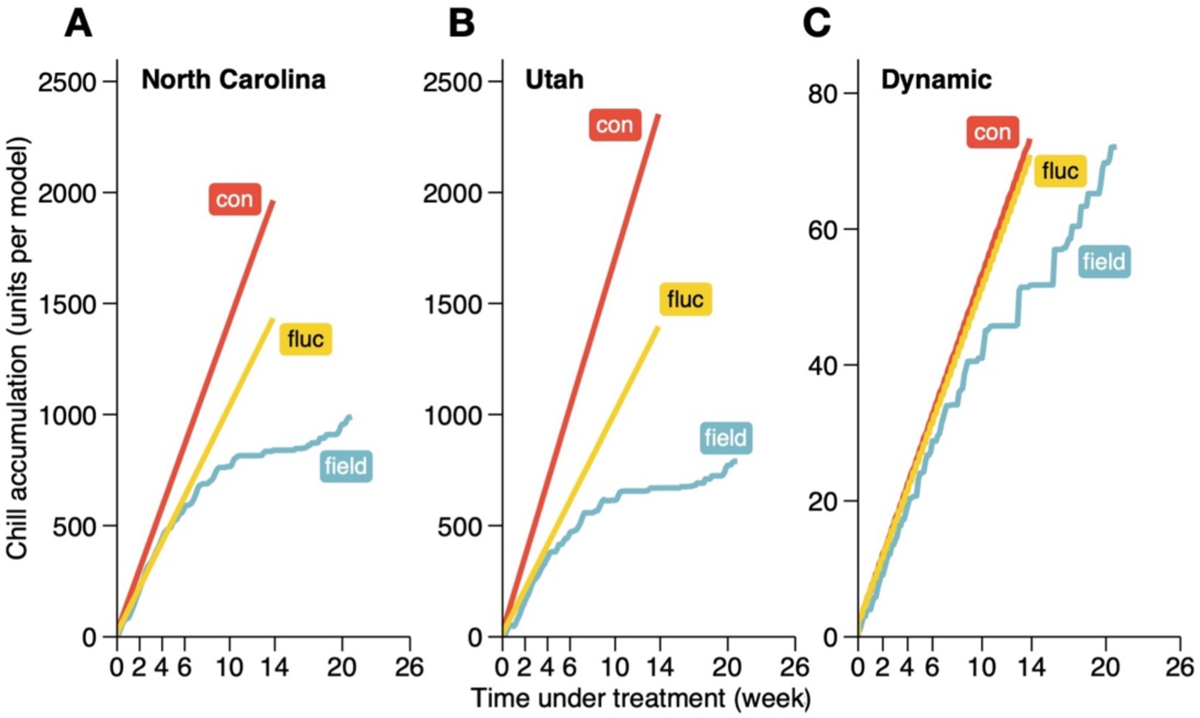
Chill accumulation based on different chill models for three chill treatments. Three temperature treatments applied to buds of interspecific hybrid grapevines (*Vitis* hybrid): constant (“con”, 5°C), fluctuating (“fluc” −3.5 to 6.5°C daily), and field conditions in Madison, WI, USA in season 2021-22. Chill accumulation was calculated for duration of treatments based on three classic models: **(A)** North Carolina (Shaltout and Unrath, 1983), **(B)** Utah (Richardson et al., 1974), and **(B)** Dynamic (Fishman et al., 1987a, b).

### Decrease in time to budbreak is differentially affected by chill treatments

At the beginning of the experiment, the average time to budbreak was 52 days (pooled for all treatments; Fig. 5A). As the duration of chill treatments increased, time to budbreak decreased towards an apparent asymptote of about 20 days to budbreak between 10 and 20 weeks of chilling exposure. Decline in time to budbreak is slightly greater in constant treatment compared to field and fluctuating temperature treatments. In the field treatment, time to budbreak started decreasing again after ∼20 weeks under treatment.

**Figure 5.**
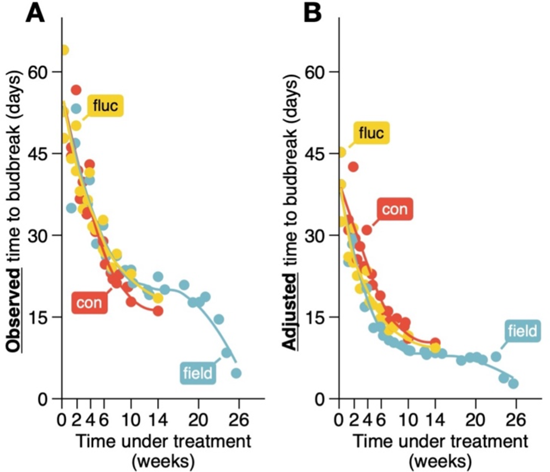
Effect of increasing time under chill treatment exposure on time to budbreak. **(A)** Time to budbreak in interspecific hybrid grapevines (*Vitis* hybrid) decreases with increased exposure to chill treatments. **(B)** Adjusted time to budbreak based on cold hardiness gains experienced during chilling treatments (Eq. 1). Chill treatments applied are: constant (“con” 5°C), fluctuating (“fluc”, −3.5 to 6.5°C daily), and field conditions in Madison, WI, USA for three seasons (2018-2019, 2019-2020, 2021-2022). Sample points indicate averages within treatments for each season and chilling duration. Lines are loess fits for each treatment in response to time.

Time to budbreak was adjusted to compensate for the gains in cold hardiness elicited by chilling treatments (Fig. 5B). The adjusted budbreak values are lower than the corresponding observed budbreak. The adjustments are greater in the field treatment compared to artificial treatments due to the greater changes in cold hardiness in the field (Fig. 3, differences along y-axis between Fig. 5A and 5B). In the spring, however, the absolute values of the adjustments approach zero for the field treatment – when deacclimation makes CH_initial_ ≅ CH_min_ and time to budbreak is short (i.e., increase in the denominator in Eq. 1). Overall, a faster decrease in adjusted time to budbreak is occurs in the field treatment, followed by fluctuating, and lastly constant treatment.

The accuracy of models was then evaluated (see Fig. 1) based on both observed and adjusted time to budbreak separately and using an exponentially declining response of time to budbreak to chill accumulation. The three classic chill models (NC, UT, and DM) had varied performances in describing chill accumulation across treatments (Fig. 6). Notably, the lower accumulation of chilling in the field treatment and greater accumulation of chill in constant treatment (see Fig. 4) led to further separation of datapoints using the NC and UT models’ chill units (thermal time) than with time (large RMSEs for field and constant, small RMSE for fluctuating). The Dynamic model was the most effective in removing the treatment structure in time to budbreak (i.e., similar RMSE across treatments and pooled).

**Figure 6.**
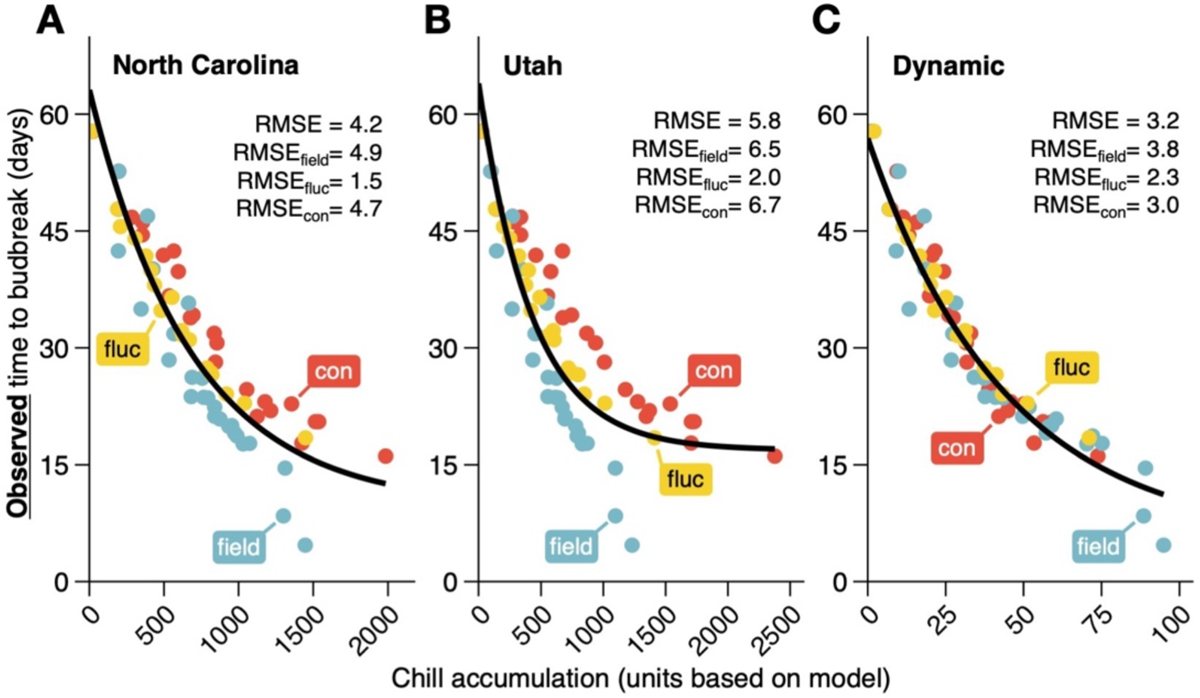
Effect of chill accumulation on observed time to budbreak. Time to budbreak of interspecific hybrid grapevine (*Vitis* hybrid) buds under forcing conditions (22 °C, 16h daylength), following pre-exposure to three different chill treatments for multiple durations. Chill treatments applied are: constant (“con”, 5 °C), fluctuating (“fluc”, −3.5 to 6.5 °C daily), and field conditions (ambient temperature in Madison, WI, USA), for three seasons (2018-2019, 2019-2020, 2021-2022). Chill accumulation was calculated based on three classic models: **(A)** North Carolina (Shaltout and Unrath, 1983), **(B)** Utah (Richardson et al., 1974), and **(B)** Dynamic (Fishman et al., 1987a, b). Sample points indicate averages within treatments for each season and chilling duration. Lines are negative exponential decline curves (Eq. 5) based on Cannell and Smith (1983), fit for all treatments combined. RMSE calculated for all treatments combined, and for each individual treatment based on residuals for the fitted curve of all treatments combined.

Regardless of which chill model is used, adjusted time to budbreak decreased faster in the field treatment compared to both artificial chill treatments (Fig. 7). Differences among treatments appeared in all three models, where RMSE is lower within the fluctuating treatment, compared to constant and field. Adjusted budbreak response differences between chill treatments still followed the same ranking order for each classic chill model: DM had the least difference among chill treatments, followed by the NC, and lastly UT.

**Figure 7.**
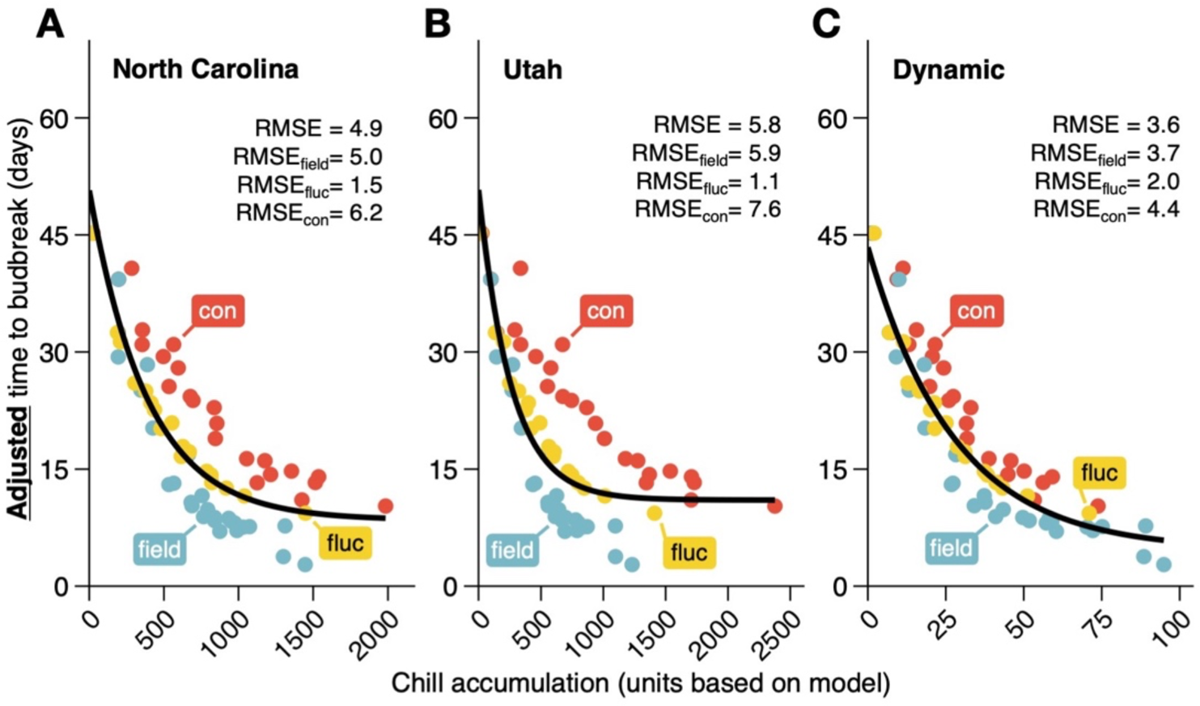
Effect of chill accumulation on cold hardiness-adjusted time to budbreak. Time to budbreak of interspecific hybrid grapevine (*Vitis* hybrid) buds under forcing conditions (22 °C, 16h daylength), following pre-exposure to three different chill treatments for multiple durations, and adjusted based on cold hardiness measured at the beginning of each budbreak assay for each treatment (Eq. 1). Chill treatments applied are: constant (“con”, 5 °C), fluctuating (“fluc”, −3.5 to 6.5 °C daily), and field conditions (ambient temperature in Madison, WI, USA), for three seasons (2018-2019, 2019-2020, 2021-2022). Chill accumulation was calculated based on three classic models: **(A)** North Carolina (Shaltout and Unrath, 1983), **(B)** Utah (Richardson et al., 1974), and **(B)** Dynamic (Fishman et al., 1987a, b). Sample points indicate averages within treatments for each season and chilling duration. Lines are negative exponential decline curves (Eq. 5) based on Cannell and Smith (1983), fit for all treatments combined. RMSE calculated for all treatments combined, and for each individual treatment based on residuals for the fitted curve of all treatments combined.

### A new model for chill accumulation

The lack of fit for adjusted days to budbreak among treatments and the incorrect chill accumulation rank for treatments according to chill accumulated in the classic models suggested the need for a new chill model. The new model was optimized based on two functions, one related to hourly temperature and the other to daily temperature amplitude. The new model has its highest efficiency at 13 °C (T_chill.opt_ in Eq. 2), which is higher than the optimum around 7.2 °C used by the classic models (Fig. 8A). Similar to classic models, the new model ceases chill accumulation at temperatures above 16 °C (T_chill.max_), but does not include chill negation as in NC and UT. Chill efficiency at 0 °C is very similar for the New model as for NC and DM. However, the low rate of decrease in efficiency allows for positive effects, albeit relatively small, in chill accumulation at negative temperatures beyond −10 °C. The enhancement effect was optimized at a maximum of 1.5-fold the efficiency (Fig. 8B). The large slope obtained for Eq. 3 results in approximately no enhancement when daily amplitude is below 3 °C, a 1.25-fold increase at 4 °C, and a maximum enhancement effect of 1.5-fold at or above 5 °C of daily amplitude.

**Figure 8.**
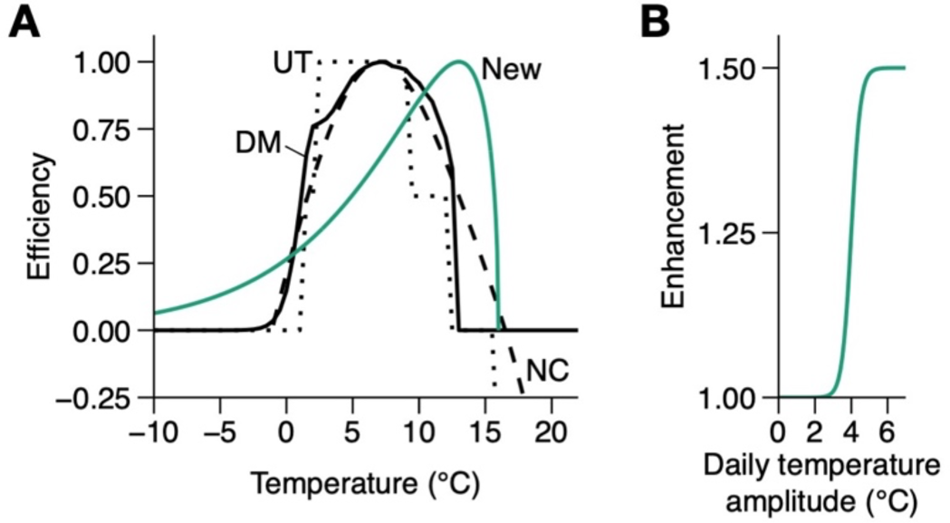
Efficiency of chill accumulation as a function of temperature for different chill models. **(A)** Efficiency of temperatures in accumulating chill for each of three classic models [North Carolina – “NC” (Shaltout and Unrath, 1983), Utah – “UT” (Richardson et al., 1974), and Dynamic – “DM” (Fishman et al., 1987a, b) models], and a newly proposed model (“New”). For DM, efficiency is calculated based on 1200 hours of constant exposure [as in Fishman et al. (1987)]. For the New model, Efficiency is calculated based on Eq. 2, where T_chill.max_ =16 °C, T_chill.opt_ =13 °C, *a = 0.5*. **(B)** Enhancement of accumulated chill for the new model as a function of daily temperature amplitude, calculated based on Eq. 3, where T_amp.infl_ = 4 °C, T_amp.slp_ = 4, *d* = 1.5.

Using the New Model, the rank order of chill accumulation changed for the different treatments. Field and fluctuating treatments alternate in terms of higher chill accumulation over time, while constant treatment generally accumulates chill at a lower rate (Fig. 9A). This change in chill accumulation inverts treatments for observed time to budbreak, where constant decreased time to budbreak with lower chill accumulated compared to field and fluctuating (Fig. 9B). However, the new model produces a better fit for observed budbreak than NC and UT models based on RMSE values. When days to budbreak are adjusted for cold hardiness, the new model removes differences in treatment response, and reduces RMSE to about half of the lowest observed in the classic models (Fig 9C).

**Figure 9.**
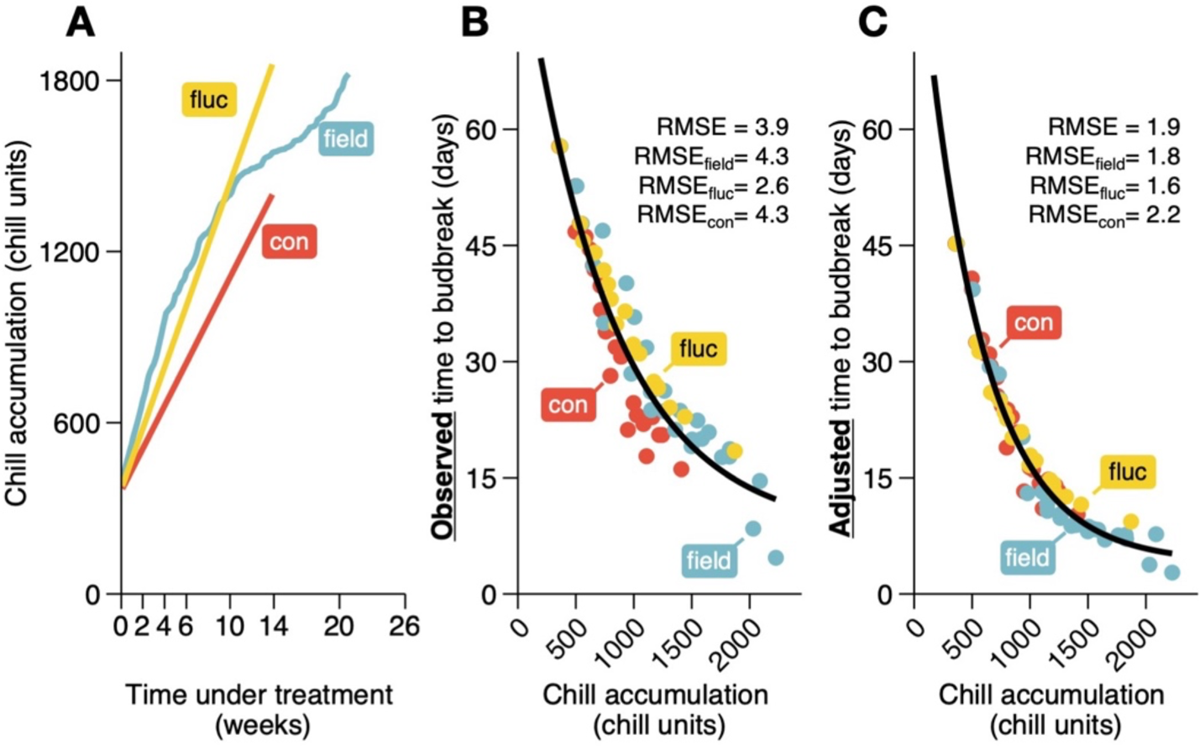
Chill accumulation and effects on time to budbreak of a new chill model. **(A)** Chill accumulation in season 2021-22 for the duration of three temperature treatments using the newly proposed chill model (Eqs. 2, 3, 4). Observed **(B)** and Adjusted time to budbreak **(C)** of interspecific hybrid grapevine (*Vitis* hybrid) buds under forcing conditions (22 °C, 16 h daylength), following pre-exposure to three different chill treatments for multiple durations as a function of chill accumulation based on the new model. Adjusted time to budbreak was calculated based on cold hardiness measured at the beginning of each budbreak assay for each treatment (Eq. 1). Chill treatments applied are: constant (“con”, 5 °C), fluctuating (“fluc”, −3.5 to 6.5 °C daily), and field conditions (ambient temperature in Madison, WI, USA), for three seasons (2018-2019, 2019-2020, 2021-2022). Sample points indicate averages within treatments for each season and chilling duration. Lines are negative exponential decline curves (Eq. 5) based on Cannell and Smith (1983), fit for all treatments combined. RMSE calculated for all treatments combined, and for each individual treatment based on residuals for the fitted curve of all treatments combined.

## Discussion

Dormancy is essential for perennial plants to survive in temperate climates and chilling is the main environmental factor involved in dormancy regulation. But the chilling efficiency of particular temperatures remains poorly understood despite extensive research in the area. To generate novel insights regarding chill accumulation, we produced the first study incorporating cold hardiness in the evaluation of effectiveness of artificial and natural chilling. The objectives in this study were (1) to evaluate the magnitude of cold hardiness changes related to chilling treatments, and (2) to incorporate cold hardiness changes in the evaluation of chill models. Our results demonstrate that cold hardiness is an important covariate in dormancy assays, where the magnitude of its effect depends on chill temperatures used. Here we highlight the importance of measuring cold hardiness, demonstrate the use of these measurements in chill accumulation evaluation, and create a new chill model framework to better describe dormancy responses observed.

### Cold hardiness evaluation is necessary to correct budbreak observations for use as a dormancy completion phenotype

Cold hardiness was differentially affected by chill treatments (Fig. 3). This reflects temperature conditions in each treatment (lower temperatures experienced = greater cold hardiness) and was expected based on models used for prediction of cold hardiness of grapevines (Ferguson et al., 2011, 2014; North et al., 2022; Kovaleski et al., 2023; Jones et al., 2023).

Concomitantly, duration of chill treatments promoted changes in dormancy status, decreasing time (under forcing) to budbreak, regardless of treatment. However, cold hardiness of buds upon entering forcing assays affects time to budbreak independently from dormancy (North and Kovaleski, 2024). Therefore, even when similar times to budbreak were observed for buds across chill treatments, there were differences in the dormancy status of buds.

The direct effect of cold hardiness on time to budbreak is simple. Buds that are hardier due to history of temperature exposure have more cold hardiness to lose before reaching the non-cold hardy budbreak stage [“path length” in Kovaleski (2022) and North and Kovaleski (2024)]. However, to deduce time to budbreak based on cold hardiness, a deacclimation rate is necessary (Kovaleski, 2022). In the present study, rather than determining deacclimation rates through repeated cold hardiness evaluation (e.g., Kovaleski et al., 2018; North et al., 2022; Kovaleski, 2022), we estimated deacclimation rates by dividing cold hardiness measured at the outset of budbreak assays by time to budbreak (denominator in Eq 1). We were able to correct budbreak observations by systematically removing uneven cold acclimation effects across chill treatments using this approach.

The correction of time to budbreak revealed issues with chilling models. Should there have been no measurements of cold hardiness, the Dynamic Model would be the best descriptor, even slightly better than the New Model (RMSE of 3.2d and 3.9d, respectively). This agrees with the observations of Luedeling and Brown (2011), where the Dynamic Model presents the best transferability across climates. It also suggests the Dynamic Model may inadvertently incorporate aspects of cold acclimation when calculating chill accumulation, but as acclimation varies depending on temperature and species (North and Kovaleski, 2024), it is possible that the Dynamic Model may fail when comparing multiple species. Most importantly, when time to budbreak was adjusted for the effects of acclimation related to treatment temperatures, clear treatment structure appeared within the Dynamic Model, even if it was still the best of the classic models (Fig. 7). Therefore, adjustments were still required in chill estimations. Rather than adapting pre-existing models, we chose to create a new framework, particularly due to effects related to temperature fluctuations.

### Distinguishing features in the new chill model

There are two notable temperature-based differences in our field and fluctuating chill conditions that are excluded from chill calculations in the existing models tested, including: (1) time at below-freezing temperatures and (2) daily temperature fluctuations. It is these differences that guided important features in our proposed new chill model.

### Freezing temperatures contribute to dormancy completion

The optimum configuration of our new chill model (lowest RMSE) included below-freezing temperatures in the temperature range that generates chill units (Fig. 8). Most existing chill models do not calculate chill units for temperatures less than 0 °C (or calculate relatively insignificant chill units for temperatures between approximately −1 and 0 °C). This may be due to lack of data from such low temperature treatments for development of models, or, when those were present, because of the competing effects of these below-freezing temperatures promoting acclimation (and thus increasing time to budbreak) and chill accumulation (and thus decreasing time to budbreak) simultaneously. Despite this confounding nature of cold hardiness effects, previous studies have still empirically demonstrated a role for temperatures less than 0 °C in dormancy completion (Mahmood et al., 2000; Rose and Cameron, 2009; Guak and Neilsen, 2013; Cragin et al., 2017; Baumgarten et al., 2021). However, the magnitude of chill accumulation effects at temperatures less than 0 °C versus traditionally accepted chill temperatures will likely have been underestimated even in these studies since cold hardiness was not evaluated simultaneously. By accounting for the effects of acclimation, we have produced a novel chill model that accumulates chill at below freezing temperatures.

### Daily temperature fluctuations contribute to dormancy completion

Temperature fluctuations have been known to contribute to chilling accumulation (Fishman et al., 1987 a, b). Yet existing chill models, except for the Dynamic Model, assume a given temperature’s contribution towards dormancy completion is constant and therefore do not account for fluctuations in temperature. This has been referred to as “time-homogeneous” (e.g., most existing chill models) versus “time-inhomogeneous” (e.g., Dynamic Model and our proposed New Model) (Zhang and Taylor, 2011). However, the combinations of temperature cycles that promote increased chill accumulation within the Dynamic Model are very specific and the magnitude of those increases are small. Neither the field nor the fluctuating chill treatments in our study accumulated more chill portions than constant when using the Dynamic Model, and therefore it could not account for the differences in adjusted time to budbreak we observed across our chill treatments.

While math and logic principles guided the equation choices for our new model, biological mechanisms were also considered. First, for the basic temperature response, an enzyme-related curve has been used to describe chill accumulation efficiency (O’Neil et al., 1972). Additionally, the enhanced chill accumulation based on daily amplitude could be an effect of factors such as entrainment of the circadian clock (Somers et al., 1998) and phytochrome effects modulated by temperature (Ding et al., 2023; Brunner et al., 2023). It is also an easy framework to reparametrize in future studies as more data becomes available. This may be particularly important as here we used *Vitis* as a model, but since species have different acclimation responses (North and Kovaleski, 2024), their responses in chill accumulation may also vary to temperatures.

### Is there a role for chill negation?

Chill negation (i.e., negative chill unit) is not present in our New Model. Both the NC and UT Models have chill negation effects at temperatures above ∼15 °C (negative efficiency in Fig. 8A). For the DM, the negation only occurs at the first step of the two-step model, where the presumed precursor of the dormancy breaking factor can be destroyed, but the process is not reversible once a unit of the dormancy breaking factor is produced. There are conflicting reports in empirical studies regarding a negative effect of warm temperatures during chill accumulation. In the studies that contributed data for the DM, some observations demonstrate a chill negation effect, but where negation only happens within the first few days of experiencing the warm temperatures (≥19 °C; Erez et al., 1979; Couvillon and Erez, 1985). However, studies of woody perennials in warmer, subtropical climates (particularly of temperate horticultural crops) have seen the need to adapt models by omitting such chill negation from warm temperatures, or accumulate chilling in a much warmer range (Ou and Chen, 2000; Lu et al., 2012; Milech et al., 2018), and the use of UT model is discouraged in warm climates (Luedeling et al., 2023). Our artificial treatments did not include temperatures where possible effects of chill negation were expected. Previous observations speculate a possible dual role of warmer temperatures in providing accumulation of both chilling and heat units, thus contributing to decreased time to budbreak (Flynn and Wolkovich, 2018). In early fall, repeated above-freezing temperature cycles have been shown to promote some acclimation (Wang et al., 2023; Hiraki et al., 2023), which could increase time to budbreak. But even low above-freezing temperatures promote loss of cold hardiness, once some chill has accumulated (Kovaleski et al., 2018; North et al., 2022; Kovaleski 2022). The effect of warm temperatures on chill negation remains unclear, but it is likely that measurements of cold hardiness of buds under different fluctuating treatments, pre-forcing, will contribute to our understanding of this effect.

## Conclusion

Here we have demonstrated a new approach for characterizing temperature-specific chill efficiency, where budbreak is still the main dormancy phenotype, but it is adjusted based on cold hardiness changes resulting from chilling treatments. Our data demonstrates that freezing temperatures do promote chilling accumulation, even if not apparent in standard budbreak responses, as they also contribute to cold acclimation. A time inhomogeneous model was necessary to describe cold hardiness-adjusted budbreak responses in this study, where we interpreted daily fluctuations as chill accumulation enhancers, as has been previously suggested. Future studies investigating chill accumulation need to include fluctuating temperatures, both experimentally and naturally derived, in their treatment designs, as well as cold hardiness measurements in order to advance realism and generality in chill models and more accurately characterize dormancy in physiological and molecular studies.

## Supporting information

Supplementary TableS1+FiguresS1-S2

## Acknowledgements

N/A

## Author Contributions

MGN, BAW, AA, and APK designed experiments; MGN collected data; MGN and APK analyzed data, prepared figures, and wrote original draft; AA and BAW contributed to writing and contextualization.

## Data Availability

We intend to make data and code available upon publication through a GitHub repository.

## Supplementary Figures

**Figure S1.** Temperatures experienced within chill treatments applied. Temperatures are shown in a grid of season by treatment. For field conditions, ambient temperature in Madison, WI, USA is shown in ribbon. For constant and fluctuating artificial treatments, field temperature is shown prior to start of experiment in mid-October of each season, and lines are used to shown max and min temperatures in each treatment.

**Figure S2.** Chill accumulation based on four different chill models for three chill treatments and three seasons. Three temperature treatments applied to buds of interspecific hybrid grapevines (Vitis hybrid): constant (“con”, 5°C), fluctuating (“fluc” −3.5 to 6.5°C daily), and field conditions in Madison, WI, USA for three seasons. Chill accumulation was calculated for duration of treatments based on three classic modelS: North Carolina (Shaltout and Unrath, 1983), Utah (Richardson et al., 1974), and Dynamic (Fishman et al., 1987a, b), and the newly proposed model.

