## Supplementary TableS1+FiguresS1-S2 for "Cold hardiness-informed budbreak reveals role of freezing temperatures and daily fluctuation in chill accumulation model"

### Contents

**Table S1.** Parameter levels tested stepwise for development of a novel chill accumulation model.
**Figure S1.** Temperatures experienced within chill treatments applied.
**Figure S2.** Chill accumulation based on four different chill models for three chill treatments and three seasons.

**Table S1.** Parameter levels tested stepwise for development of a novel chill accumulation model.

| Parameter | Units | Start | Stop | Interval |
|---|---|---|---|---|
| $T_{chill.max}$ | °C | 0 | 24 | 2 |
| $T_{chill.opt}$ | °C | -6 | 16 | 1 |
| *a* | no units | NA | NA | 0.5, 0.6, 0.7, 0.8, 0.9, 1.0, 1.25, 1.5, 2.0, 2.5, 3.0, 4.0, 5.0, 7.5, 10, 15 |
| $T_{amp.infl}$ | °C | 0 | 8 | 2 |
| $T_{amp.slp}$ | no units | NA | NA | 0.5, 1.0, 2.0, 4.0, 8.0, 16 |
| *d* | no units | 1 | 2 | 0.5 |

Combinations where $T_{chill.max} < T_{chill.opt}$ were removed (430,560 total combinations for all parameters, 103,680 removed; 313,920 combinations tested).

In **Equation 2**, $T_{chill.max}$ (°C) is the maximum temperature for chilling, $T_{chill.opt}$ (°C) is the optimum temperature for chilling, *"a"* (no units) is the coefficient related to the slope of the curve.

In **Equation 3**, $T_{amp.infl}$ (°C) is the inflection point of the curve, $T_{amp.slp}$ (no units) is the slope of the curve, "d" (no units) is the upper chill enhancement limit.

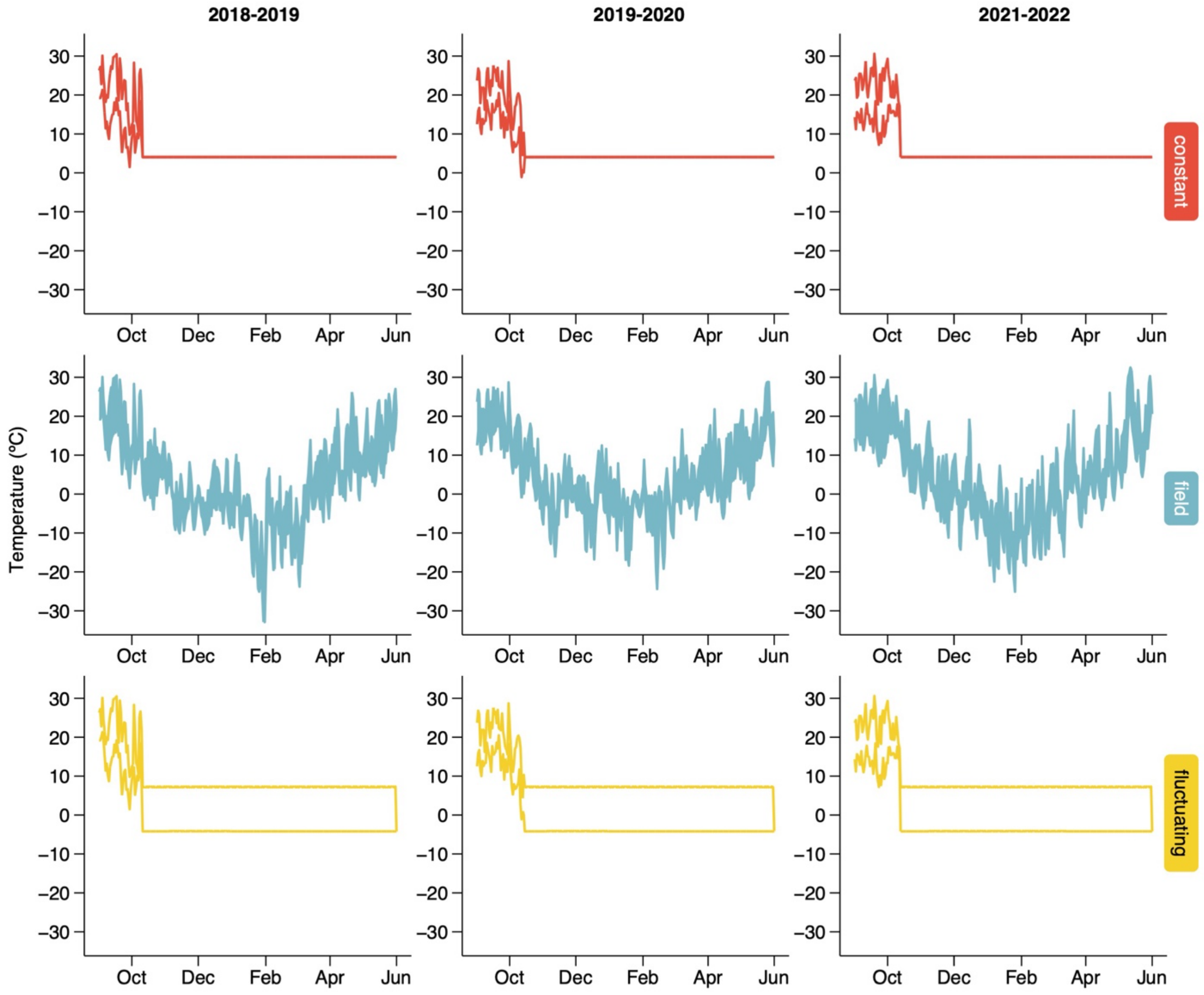


**Figure S1. Temperatures experienced within chill treatments applied.** Temperatures are shown in a grid of season by treatment. For field conditions, ambient temperature in Madison, WI, USA is shown in ribbon. For constant and fluctuating artificial treatments, field temperature is shown prior to start of experiment in mid-October of each season, and lines are used to shown max and min temperatures in each treatment.

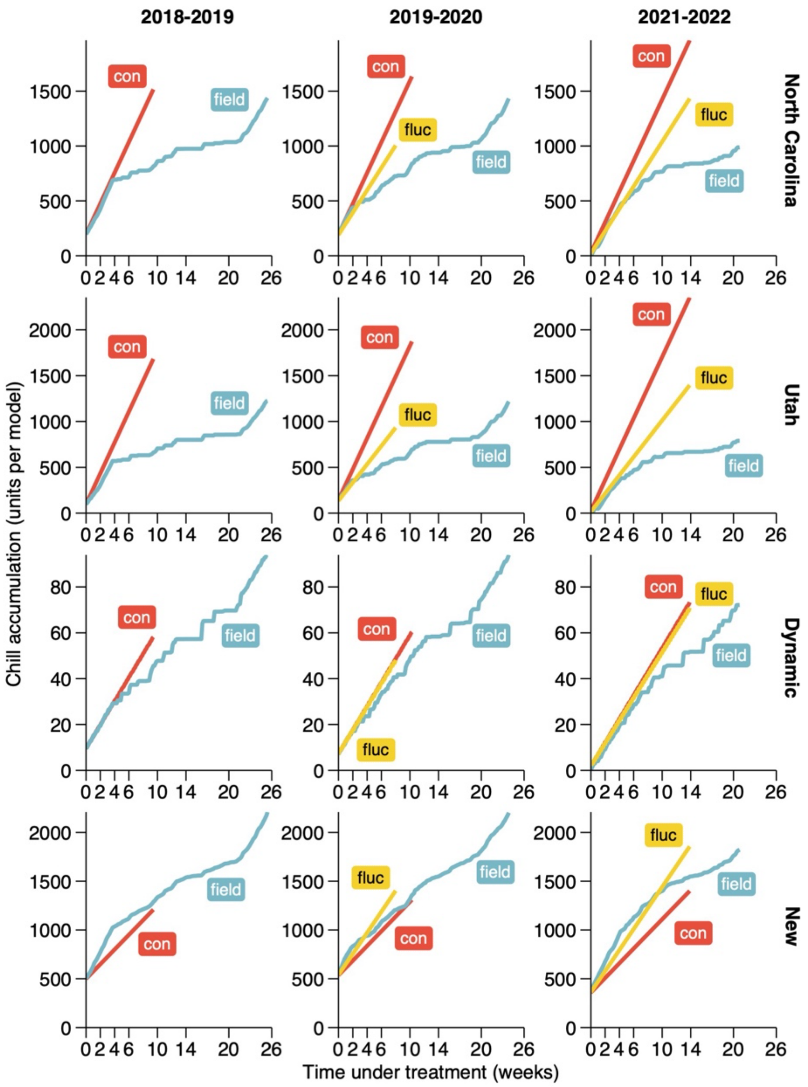


**Figure S2. Chill accumulation based on four different chill models for three chill treatments and three seasons.** Three temperature treatments applied to buds of interspecific hybrid grapevines (Vitis hybrid): constant ("con", 5°C), fluctuating ("fluc" -3.5 to 6.5°C daily), and field conditions in Madison, WI, USA for three seasons. Chill accumulation was calculated for duration of treatments based on three classic models: North Carolina (Shaltout and Unrath, 1983), Utah (Richardson et al., 1974), and Dynamic (Fishman et al., 1987a, b), and the newly proposed model.
